# The *Peroxiredoxin 6* gene plays a critical role in the homeostatic regulation of fear response

**DOI:** 10.1101/2020.07.10.196477

**Authors:** Sarayut Phasuk, Tanita Pairojana, Pavithra Suresh, Shun-Ping Huang, Narawut Pakaprot, Supin Chompoopong, Chee-Hing Yang, Hsueh-Kai Chang, Chien-Chang Chen, Ingrid Y. Liu

## Abstract

Peroxiredoxin 6 (PRDX6) is a multifunctional enzyme implicated in redox regulation and expressed in many organs including the brain. It is known to participate in many psychiatric functions, but its role in fear memory is unknown. The present study demonstrates that *PRDX6* plays a critical role in the regulation of fear response. Using *Prdx6* knockout (*Prdx6*^*−/−*^) mice, we identified that PRDX6 acts as a suppressor in fear memory formation. Lack of *Prdx6* leads to the faster fear acquisition and enhanced contextual fear response. This phenomenon was confirmed by the fact that injection of lentivirus-carried human PRDX6-V5 into the hippocampus of *Prdx6*^*−/−*^ mice restored the enhanced fear response to the wild-type level. In the hippocampus of *Prdx6*^*−/−*^ mice, calcium-dependent PLA2 level was increased, which may compensate for the lack of aiPLA2 function to maintain normal synaptic membranes. On the other hand, reactive oxygen species (ROS) levels did not change, indicating loss of peroxidase function did not affect the regulation of fear response.

## Introduction

Approximately 8-10% of those who experience traumatic events develop posttraumatic stress disorder (PTSD), and more than 50% of PTSD cases are not responsive to psychological therapies (1-4). To date, no effective treatment is available for curing PTSD; therefore, an understanding of the etiology of fear memory dysregulation is necessary for the development of efficient therapeutic strategies. A maladaptive fear response to previous traumatic stress is known to be the characteristic PTSD symptoms (5, 6), which involves in several brain regions including the amygdala, medial prefrontal cortex, and hippocampus (7-9). Peroxiredoxin 6 (PRDX6) is first described as an antioxidant enzyme that involves in oxidative homeostasis and lipid turnover (. PRDX6 expressed throughout the abovementioned regions (10, 11) and is a multi-functional enzyme with multiple activities. With homology to peroxiredoxin family proteins, PRDX6 displays a peroxidase activity towards peroxides such as H_2_O_2_, short-chain organic fatty acids, and phospholipid hydroperoxides. PRDX6 also exhibits acidic calcium-independent phospholipase A2 (aiPLA2) and lysophosphatidylcholine acyltransferase (LPCAT) activities, which determine its roles in various organs under different physiological and pathobiological conditions (12, 13). According to a genome-wide association study (GWAS) and proteomic studies in human patients, PRDX6 expression in the brain is altered in different types of psychiatric disorders, including major depressive disorder (14), suicidal behavior (14), and schizophrenia (3, 15), but its role in PTSD is not yet clear. With the high comorbidity of these psychiatric disorders, it is possible that PRDX6 participates in PTSD pathological development as well (16). Several pieces of evidence have revealed a strong association of hypothalamic-pituitary-adrenal (HPA) axis activity with PTSD pathology. In addition, glucocorticoid has been identified to be an important effector in HPA axis activity and fear memory formation (17, 18). Since PRDX6 is positively regulated by dexamethasone, a glucocorticoid (GC) homolog, it may act as a stress-responsive protein and affect fear memory formation (19). Additionally, PRDX6 can modulate several signaling molecules tightly associated with an abnormal fear response; thus, we hypothesize that PRDX6 may play an important role in the processing of fear memory (20-23). In the present study, we demonstrate a series of evidences to prove that *Prdx6* gene plays a critical role in the regulation of traumatic fear response. Since there is no report about PRDX6 expression in PTSD pathology, we first observed the expression levels of total PRDX6 protein in the hippocampi of wild-type mice subjected to acute restraint stress and fear conditioning, mouse paradigms mimicking PTSD. We examined the relationship between PRDX6 expression and glucocorticoid (GC), a stress hormone by treating retinal pigment epithelium (RPE) cells with GC. To investigated the role of PRDX6 underlying trace fear conditioning, we performed a loss-of-function study by employing *Prdx6*^*−/−*^ mice in trace fear conditioning (TFC) experiments. To further confirm the significance of PRDX6 in processing fear memory, we injected lentivirus-carrying human PRDX6 (hPRDX6) with a V5 tag (LV-hPRDX6-V5) into the 3^rd^ ventricle for gain-of-function experiments. Cellular and molecular studies were also conducted to go insight into the mechanisms in which lack of PRDX6 caused contextual memory enhancement. Finally, we investigated cell types and the distribution of PRDX6 expression in the hippocampus after fear memory retrieval.

## Results

### PRDX6 expression is reduced under traumatic stress

To understand whether the PRDX6 protein responds to traumatic stress, we first investigated the expression of PRDX6 after two types of stressful training-trace fear conditioning (TFC) and acute restraint stress (RST) both are well accepted as PTSD models (24). Next, C57BL/6J mice were subjected to TFC (Fig. 1A). Results showed that the baselines of freezing percentage for naive and trained mice were similar (Fig. 1B) (*t*_18_ = – 0.375, *p* = 0.712). After the first tone and shock pair, trained mice exhibited significantly increased freezing percentages during 3 trials (Fig. 1B) [trial 1: *F*_1,18_ = 14.342; *p* = 0.001; trial 2: *F*_1,18_ = 17.018; *p* = 0.001; trial 3: *F*_1,18_ = 28.781; *p* = 0.000]. The total freezing percentage (learning ability) of the trained group was significantly higher than that of the naive group (Fig. 1C) (*t*_17_ = −6.411, *p* = 0.000). Trained mice also displayed typically higher fear memory retrieval to context than the naive group (Fig. 1D) (*t*_16_ = −3.563, *p* = 0.003). We next measured PRDX6 protein expression in the hippocampus during memory consolidation (3 hours after training) and retrieval (20 minutes after the contextual test) stages (Fig. 1E, F; upper panel). During both stages, the total PRDX6 protein expression was significantly decreased (Consolidation stage, Fig. 1E, *p* = 0.023), (Retrieval stage, Fig. 1F, *p* = 0.033). The C57BL/6J mice were immobilized for 30 minutes in a 50 ml cylinder tube to induce RST and were sacrificed immediately after completion (Fig. 1G; upper panel). RST mice exhibited significantly lower total PRDX6 in the hippocampus than the non-RST group (Fig. 1G; lower panel and H) (*t*_6_ = 3.449, *p* = 0.014). Both RST and TFC enhanced plasma GC release (17), and PRDX6 can be regulated by dexamethasone-a GC homolog (19). To investigate PRDX6 expression level under GC treatment, we applied different doses of GCs (1, 20, and 100 nM) to human retinal pigment epithelium (RPE) cells that can express both PRDX6 and steroid receptors. MTT assay showed that GC treatment at all doses did not cause cytotoxic effect (Fig. S2A) [*F*_4,25_ = 0.191; *p* = 0.941]. Subsequent immunocytochemistry staining revealed that PRDX6 expression was significantly reduced under 100 nM GC treatment (Fig. 1I and J) [*F*_4,25_ = 4.506; *p* = 0.008].

**Figure 1.**
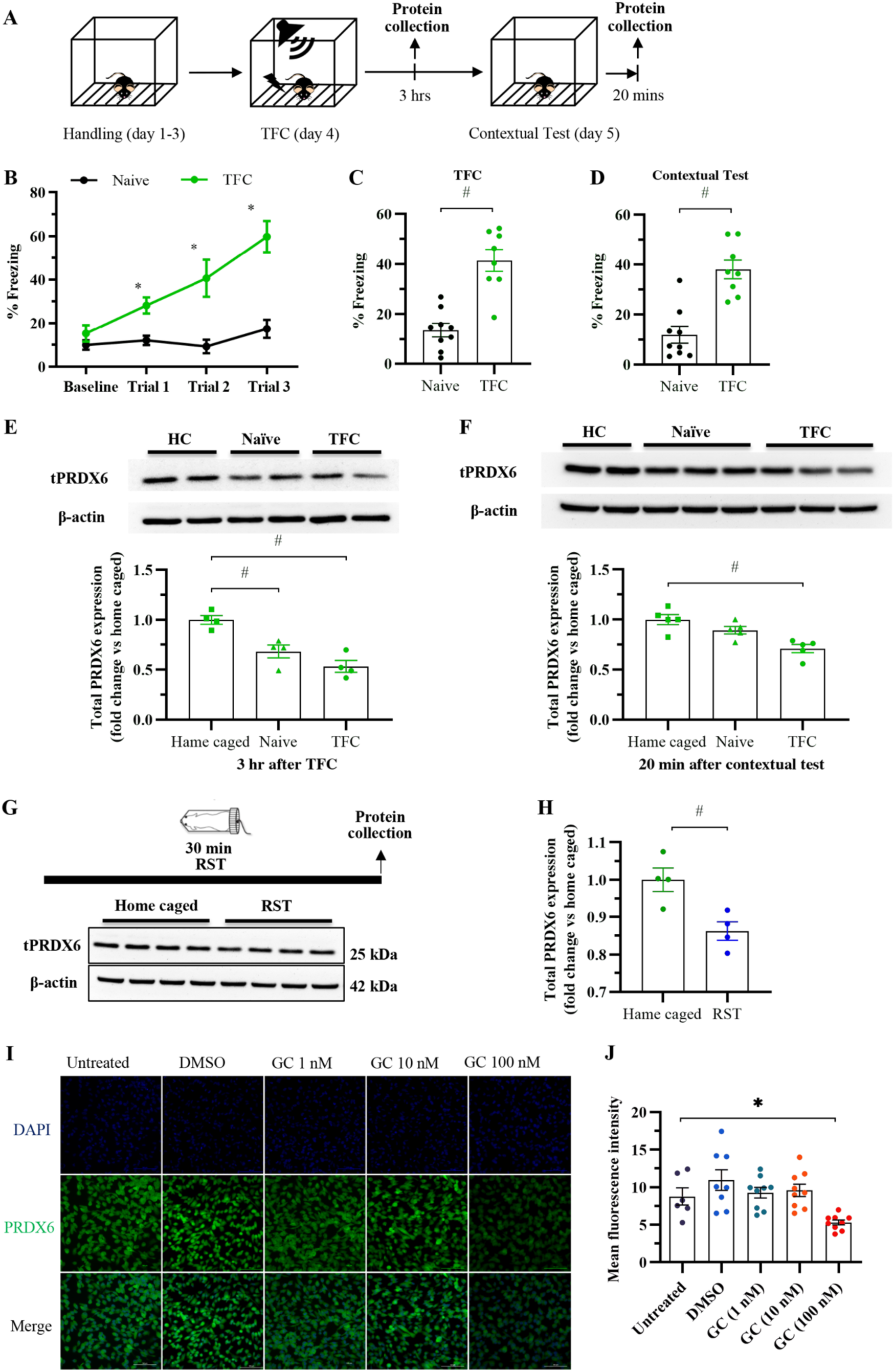
Glucocorticoid may regulate total PRDX6 level in response to different PTSD models. **(A)** Hippocampal protein samples were collected 3 hours after the TFC and 20 minutes after the contextual test. **(B)** Learning curve of baseline and after each tone-shock pair (n = 8-9 mice per group). **(C)** Total freezing percentage during the training session. **(D)** Freezing percentage during the contextual test of mice. **(E; upper panel)** Immunoblotting of the expression of total PRDX6 in the hippocampus. **(E; lower panel)** Quantification data for the expression of total PRDX6 in the hippocampus 3 hours after training (one-way analysis of variance (ANOVA) with LSD’s post hoc test). The analysis revealed a decrease in total PRDX6 during memory consolidation. **(F; upper panel)** Immunoblotting of the expression of total PRDX6 in the hippocampus. **(F; lower panel)** Quantification data for the expression of total PRDX6 in the hippocampus 20 minutes after the contextual test. The analysis revealed a decrease in total PRDX6 during memory consolidation and retrieval. **(G; upper panel)** Hippocampal proteins were collected immediately after the completion of immobilization. **(G; lower panel)** Immunoblots of the expression of total PRDX6 and β-actin in the hippocampus (n = 4 per group). **(H)** Quantification data for the expression of total PRDX6 (Student’s *t* test) in the hippocampus as a result of acute restraint stress. Suppression of PRDX6 expression by glucocorticoid (GC) treatment in RPE cells. **(I)** Immunofluorescent images of PRDX6 expressed in RPE cells after GC treatment. **(J)** Quantification of PRDX expression in response to GC treatment. All data represent the mean ± the SEM. \**p* < 0.05. Acute RST, acute restraint stress; PRDX6, peroxiredoxin 6; HC, home caged; TFC, trace fear conditioning; GC, glucocorticoid.

### *Prdx6*^*−/−*^ mice exhibited enhanced contextual and tone memory to trace fear conditioning

To identify the function of PRDX6 in the regulation of the fear response, *Prdx6*^*−/−*^ mice first underwent trace fear conditioning (TFC). The protocol used for TFC is schemed in Fig. 2A. During the first three days, mice were placed in the conditioning chamber and acclimatized to the context for 15 minutes per day. On day 4, TFC was applied, followed by contextual test 24 h later. No significant difference in learning ability (Fig. 2B) [*F*_3,27_ = 2.349; *p* = 0.095] and total freezing percentage during TFC training (Fig. 2C) (*t*_18_ = −0.947, *p* = 0.356) between the *Prdx6*^+*/*+^ and *Prdx6*^*−/−*^ mice. Interestingly, the *Prdx6*^*−/−*^ mice exhibited a significantly higher freezing percentage to context (Fig. 2D) (*t*_18_ = −4.911, *p* = 0.000) than the *Prdx6*^+*/*+^ mice. The *Prdx6*^*−/−*^ mice exhibited hyperlocomotion compared with the *Prdx6*^+*/*+^ control group, therefore, higher freezing response to context did not result from abnormal locomotion. The total distance traveled (Fig. 2E) (*t*_38_ = −2.665, *p* = 0.011) and moving speed (Fig. 2F) (*t*_38_ = −2.667, *p* = 0.011) of the *Prdx6*^*−/−*^ mice were significantly higher than those of the *Prdx6*^+*/*+^ mice. The *Prdx6*^+*/*+^ mice showed similar result in elevated plus maze as controls indicated by equal time spent in open arms (Fig. 2G) (*t*_36_ = −0.274, *p* = 0.785) and close arms (Fig. 2H) (*t*_36_ = −0.180, *p* = 0.858). To confirm the role of PRDX6 in the regulation of fear memory, gain-of-function study was conducted by intracerebroventricularly injecting lentivirus-carrying hPRDX6-V5 into the third ventricle near the hippocampal region of *Prdx6*^*−/−*^ mice. Figure 3A and 3B illustrate the site of injection and lentiviral construct. The *Prdx6*^*−/−*^ mice were intracerebroventricularly injected with either LV-hPRDX6 or LV-EGFP (7 ×10^5^ TU in a total volume of 2 μl) and then allowed to recover for 2 weeks before being subjected to TFC (Figure 3C). Fluorescent images demonstrated the expression of EGFP in three hippocampal regions, including CA1, CA3 and DG (Fig. 3D). The learning curve (Fig. 3E) and the total freezing percentage during the training session were similar between the two groups (Fig. 3F) (*t*_6_ = 1.521, *p* = 0.179), indicating that the injection of hPRDX6-V5 did not affect the learning ability of the *Prdx6*^*−/−*^ mice. Lentiviral injection of hPRDX6-V5 successfully reversed the *Prdx6* knockout effect on contextual fear memory (Fig. 3G) (*t*_6_ = 2.521, *p* = 0.045). This result suggests that hippocampal PRDX6 is involved in regulating memory retrieval to the context.

**Figure 2.**
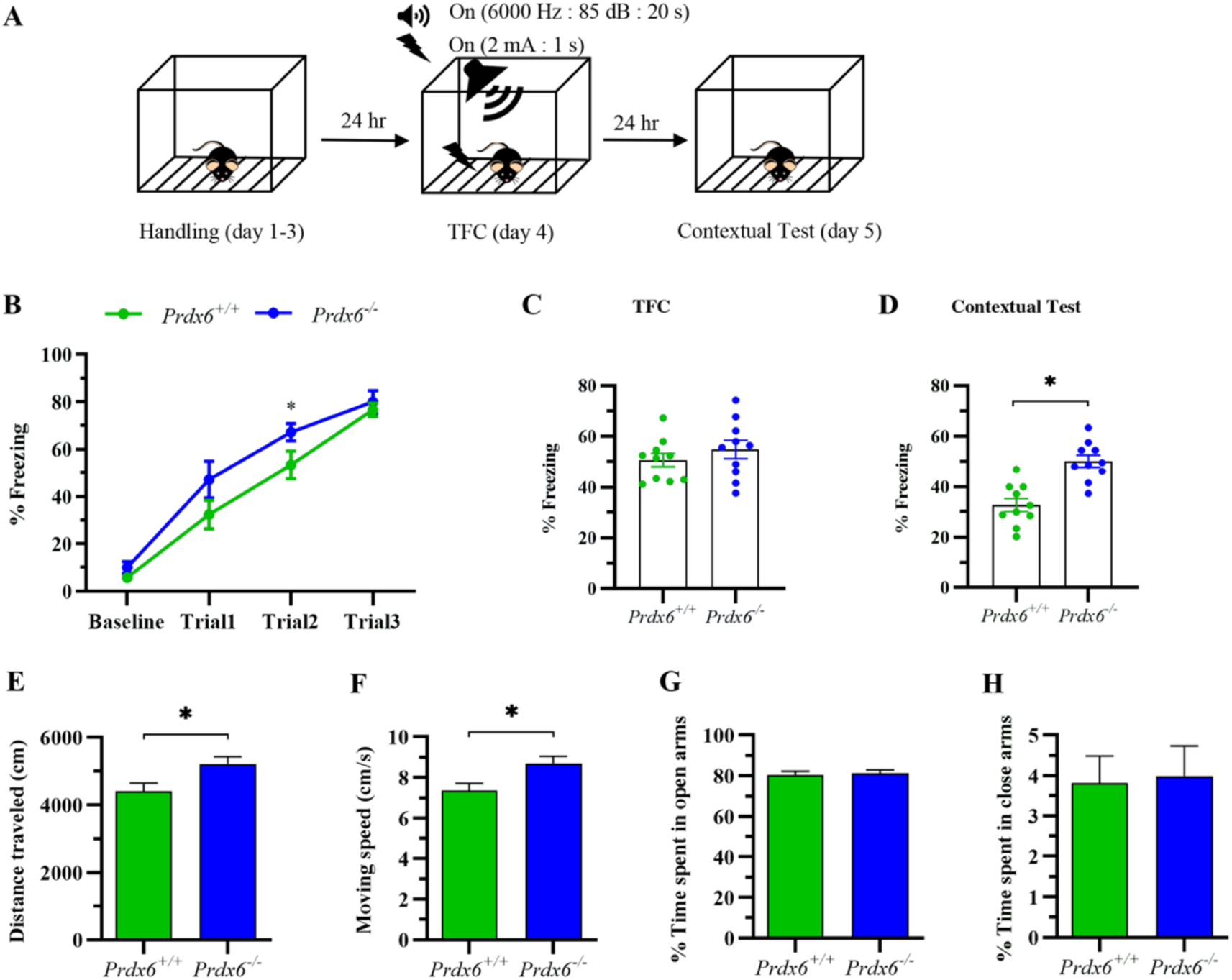
Loss of the *Prdx6* gene caused an enhanced fear response, which was not due to abnormal of neither locomotion nor anxiety. **(A)** General procedure for trace fear conditioning: Habituation to the chamber was performed for 3 consecutive days. The next day, mice were conditioned with three tone and shock pairs. Contextual fear memories were tested 24 hours later followed by a tone test to evaluate cue fear memory. **(B)** Learning curve of baseline after each tone-shock pair with a statistically significant difference at trial 2 (Student’s *t* test). **(C)** Total freezing percentage of *Prdx6*^+/+^ or *Prdx6*^−/−^ mice (n = 10/group) during the training session. **(D)** Total freezing percentage during the contextual test of mice. **(E)** Quantification data of distance traveled for 10 minutes (n = 19-20/group, Student’s *t* test). (**F)** The mean moving speed (cm/s) of the mice introduced in the open field test. **(G**) Percent time spent in open arms (n = 18-20/group). **(H)** Percent time spent in close arms. All data represent the mean ± the SEM. \**p* < 0.05. TFC, trace fear conditioning.

**Figure 3.**
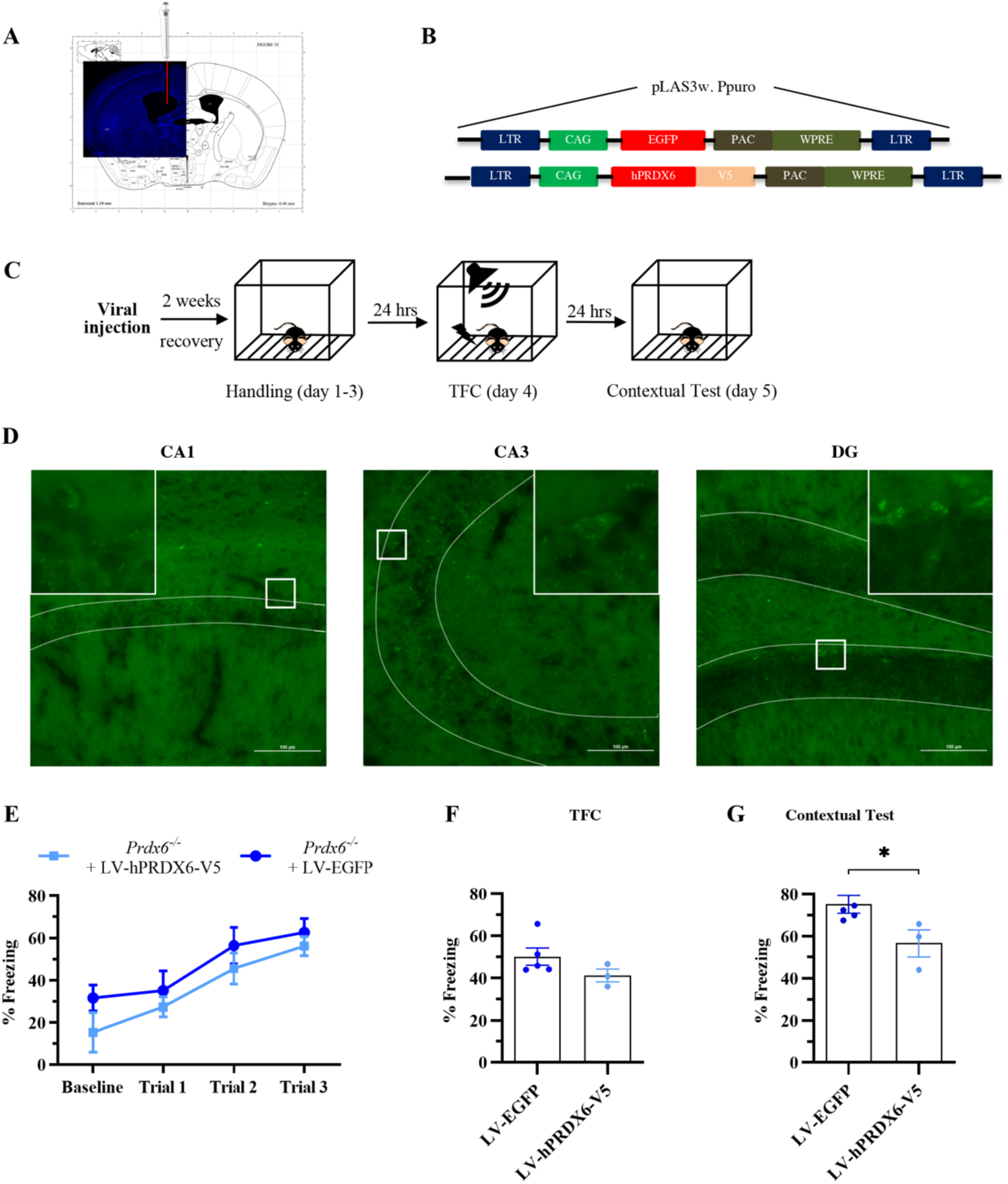
Enhanced fear memory in *Prdx6*^−/−^ mice was rescued by the injection of LV-hPRDX6-V5 into the lateral third ventricle. **(A)** Representative image of cannula tip position (red) in the right lateral ventricle. **(B)** The schematic lentivirus construct pLAS3w. *Ppuro* contains either EGFP or hPRDX6-V5. Expression of EGFP across the different subregions in the hippocampus. **(C)** Fluorescent images of EGFP expression in the CA1, CA3 and DG subregions of the hippocampus 2 weeks after intraventricular injection. **(D)** The procedure for the overexpression study. Mice were injected with lentivirus and housed for 2 weeks before performing trace fear conditioning. **(E)** Learning curve of baseline and each tone-shock pair (one-way repeated-measure ANOVA with Bonferroni’s post hoc test. **(F)** Total freezing percentage of *Prdx6*^*−/−*^ with EGFP (n = 5) or *Prdx6*^*−/−*^ with hPRDX6-V5 (n = 3) mice during the training session. **(G)** Freezing percentage during the contextual test of mice. All data represent the mean ± the SEM. \**p* < 0.05. RST, restraint stress. TFC, trace fear conditioning; EGFP, enhanced green fluorescent protein; hPRDX6-V5, human PRDX6-V5.

### Reactive oxygen species (ROS) level in the hippocampus of *Prdx6*^*−/−*^ mice after contextual test was the same as controls

Several studies have revealed the antioxidative effect of PRDX6 through its peroxidase activity in many organs including the brain (25, 26). In addition, the alteration of ROS level significantly influences the processes of memory which can be suppressed or enhanced depending on the levels of ROS (27, 28). To determine whether lack of PRDX6 alters the superoxide level in the hippocampus, we performed dihydroethidium (DHE) staining-a superoxide-sensitive dye staining. The brains were collected 20 mins after contextual test (Fig. 4A). Figure 4B shows the ethidium fluorescence of DHE. Quantitative analysis showed no significant difference in the DHE-positive density was found between genotypes in both the CA1 (Fig. 4C) (*t*_4_ = −0.508, *p* = 0.638) and CA3 (Fig. 4D) (*t*_4_ = −0.060, *p* = 0.0.955) subregions of the hippocampus.

**Figure 4.**
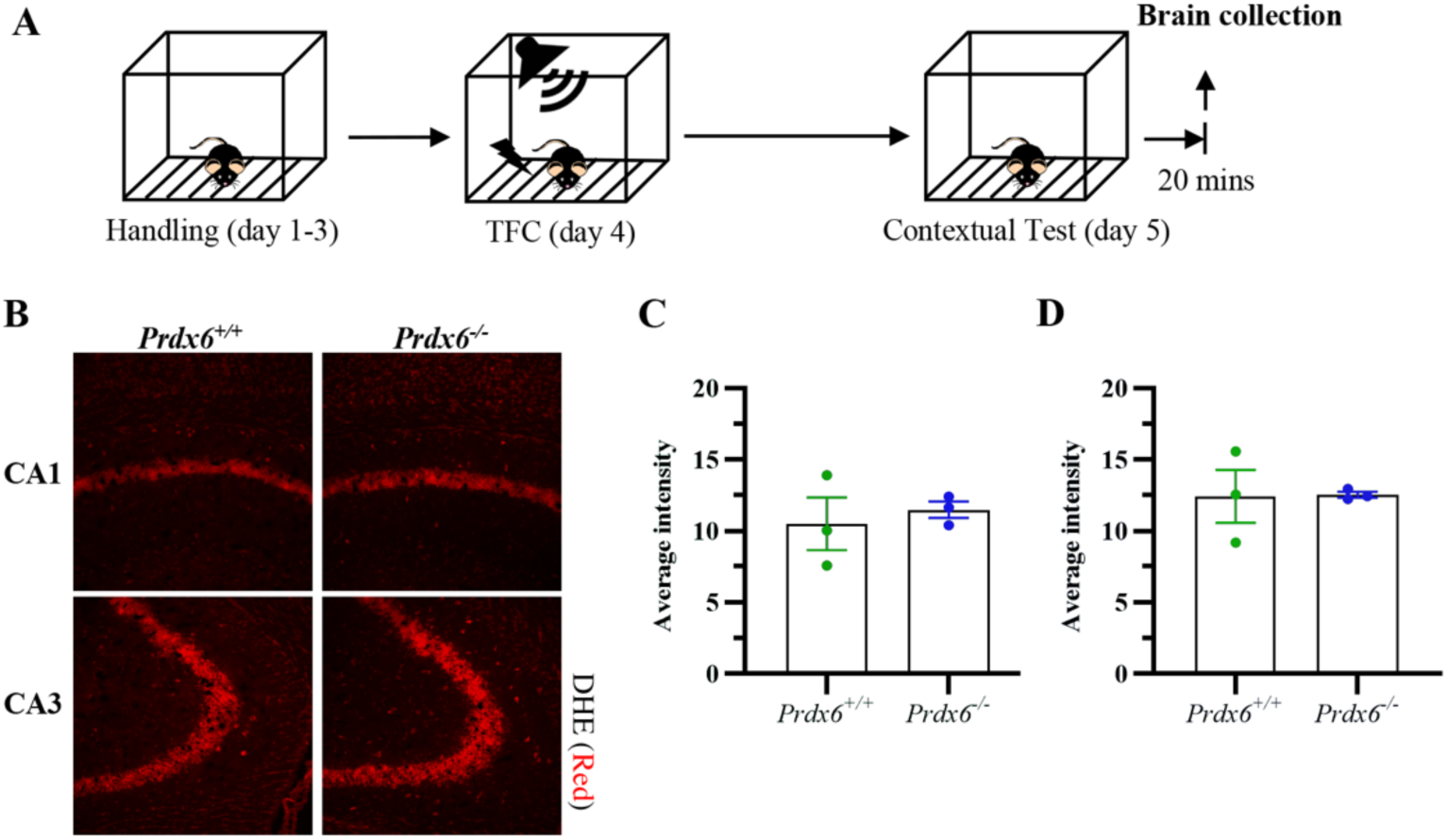
Superoxide-sensitive DHE staining in hippocampal CA1 and CA3 regions of *Prdx6*^+*/*+^ and *Prdx6*^*−/−*^ mice. **(A)** Brains were isolated 20 minutes after the contextual test. ROS levels in subfields of the hippocampus after memory reactivation remain the same. **(B)** Fluorescence images of the hippocampal subregions, including CA1 and CA3, were stained with DHE. **(C and D)** Quantification data showing similar ROS levels in the hippocampal CA1 (n = 3/group) and CA3 of *Prdx6*^*−/−*^ mice in comparison with *Prdx6*^+*/*+^ mice, respectively. All data represent the mean ± the SEM. \**p* < 0.05. CA1, cornu ammonis 1; CA3, cornu ammonis 3; DG, dentate gyrus.

### Late-phase of hippocampal LTP was reduced in *Prdx6*^*−/−*^ mice

To understand the cellular mechanism underlying the enhanced expression of fear memory in the *Prdx6*^*−/−*^ mice, we then conducted LTP recording in the CA1 region of acute hippocampal slices. The enhancement of LTP magnitude averaged throughout 3 hours after high-frequency stimulation (HFS) was seen in both *Prdx6*^+*/*+^ and *Prdx6*^*−/−*^ mice (Fig. 5A) (*t*_4_ = −13.935, *p* = 0.000 and *t*_4_ = −10.666, *p* = 0.000, respectively). In addition, the average percentage of baseline fEPSP slope evoked in Schaffer collateral stimulation was equal in both groups (Fig. 5B) (*t*_8_ = 0.59, *p* = 0.571). Early phase of LTP was similar (during 0 min to 60 min after HFS) (Fig. 5C) (*t*_8_ = 0.879, *p* = 0.405) between genotypes. Unexpectedly, we found decrease in LTP slope started from 2 hours after HFS and was maintained at lower level until late phase of LTP (during 120 min to 180 min after HFS) (Fig. 5D) (*t*_8_ = 4.005, *p* = 0.004) in hippocampal slices prepared from *Prdx6*^*−/−*^ mice. These results suggested that the lack of *Prdx6* causes impairment in the induction and the maintenance of late-LTP in the hippocampal CA1 region, which was contradict to typical reports that memory enhancement is associated with increased late-phase LTP.

**Figure 5.**
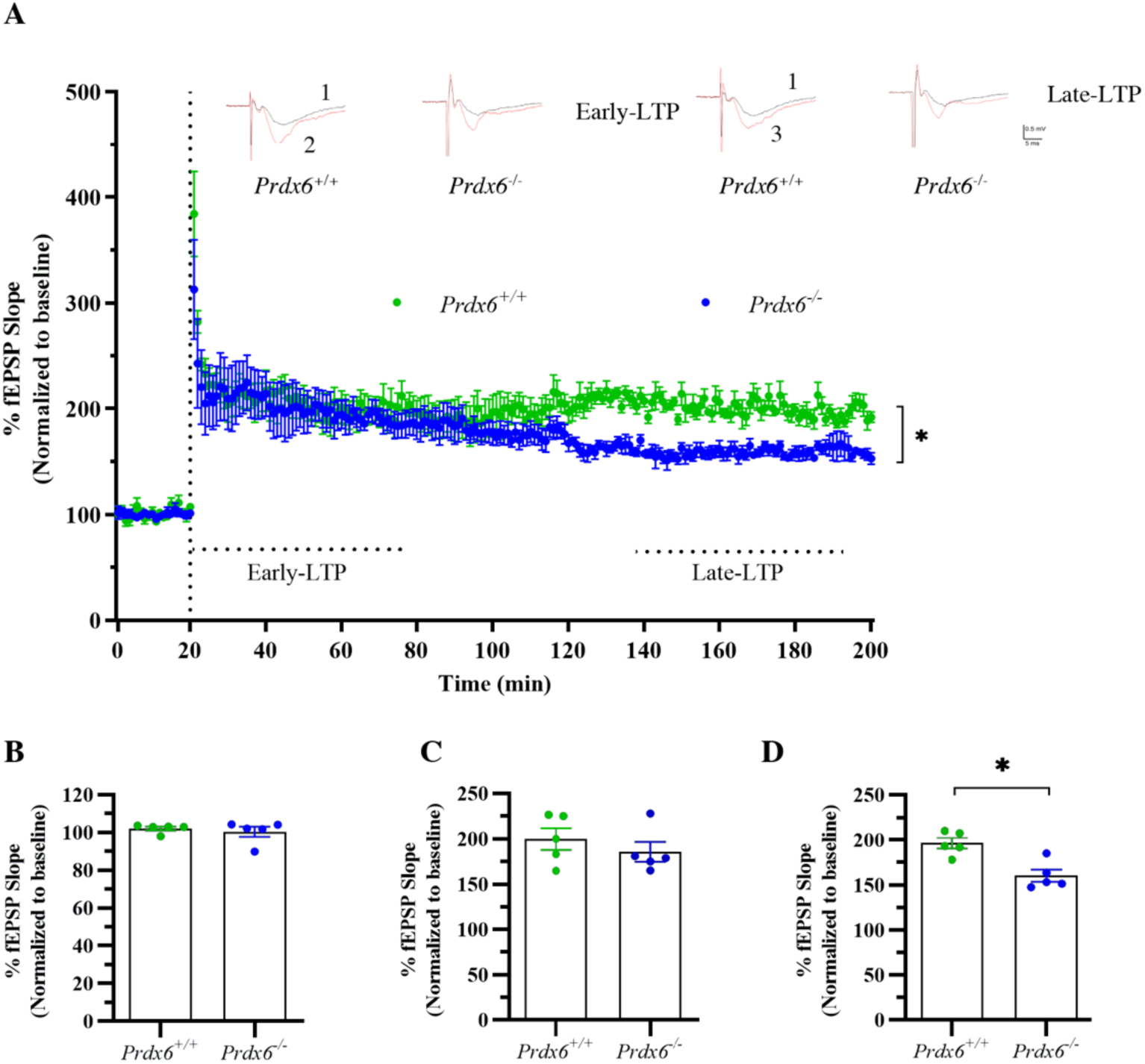
Late-phase long-term potentiation (LTP) was impaired in the hippocampi of *Prdx6*^−/−^ mice. **(A)** Summary plots of normalized LTP slope (%) before and after high-frequency stimulation (HFS) (n = 4/group, Student’s *t* test) **(B)** Normalized fEPSP amplitude (%) for baseline. **(C)** Normalized fEPSP amplitude (%) for early-LTP (last 10 minutes of first hour after HFS). **(D)** Normalized fEPSP amplitude (%) for late-LTP (last 10 minutes of the third hours after HFS). All data represent the mean ± the SEM. \**p* < 0.05.

### Expression levels of fear memory associated molecules were altered in the hippocampi of *Prdx6*^*−/−*^ mice during memory consolidation and retrieval

To understand the changes of expression levels for memory- and PRDX6-associated molecules during consolidation stage (3 hours after training) and retrieval stage (20 minutes after contextual memory test) (Fig. 6A), we measured expression levels of total cytosolic phospholipase A2 (cPLA2) and the phosphorylated extracellular signal-regulated kinases 1 and 2 (ERK1/2), both are involved in memory consolidation and retrieval; and interleukin 6 (IL-6) and its downstream molecule, Janus kinase 2 (JAK2), that are involved in memory consolidation. Western blot analysis showed that cPLA2 level was significantly elevated in the *Prdx6*^*−/−*^ mice (Fig. 6B) (*t*_5_ = −3.158, *p* = 0.025). In contrast, the IL-6 level (Fig. 6C) (*t*_5_ = 3.764, *p* = 0.013), phospho-JAK2 (Fig. 6D) (*t*_5_ = 4.248, *p* = 0.008), in *Prdx6*^*−/−*^ mice were reduced in both stages. On the other hand, phospho-ERK1/2 (Fig. 6E) (*t*_5_ = 3.638, *p* = 0.015) was reduced in memory consolidation stage.

**Figure 6.**
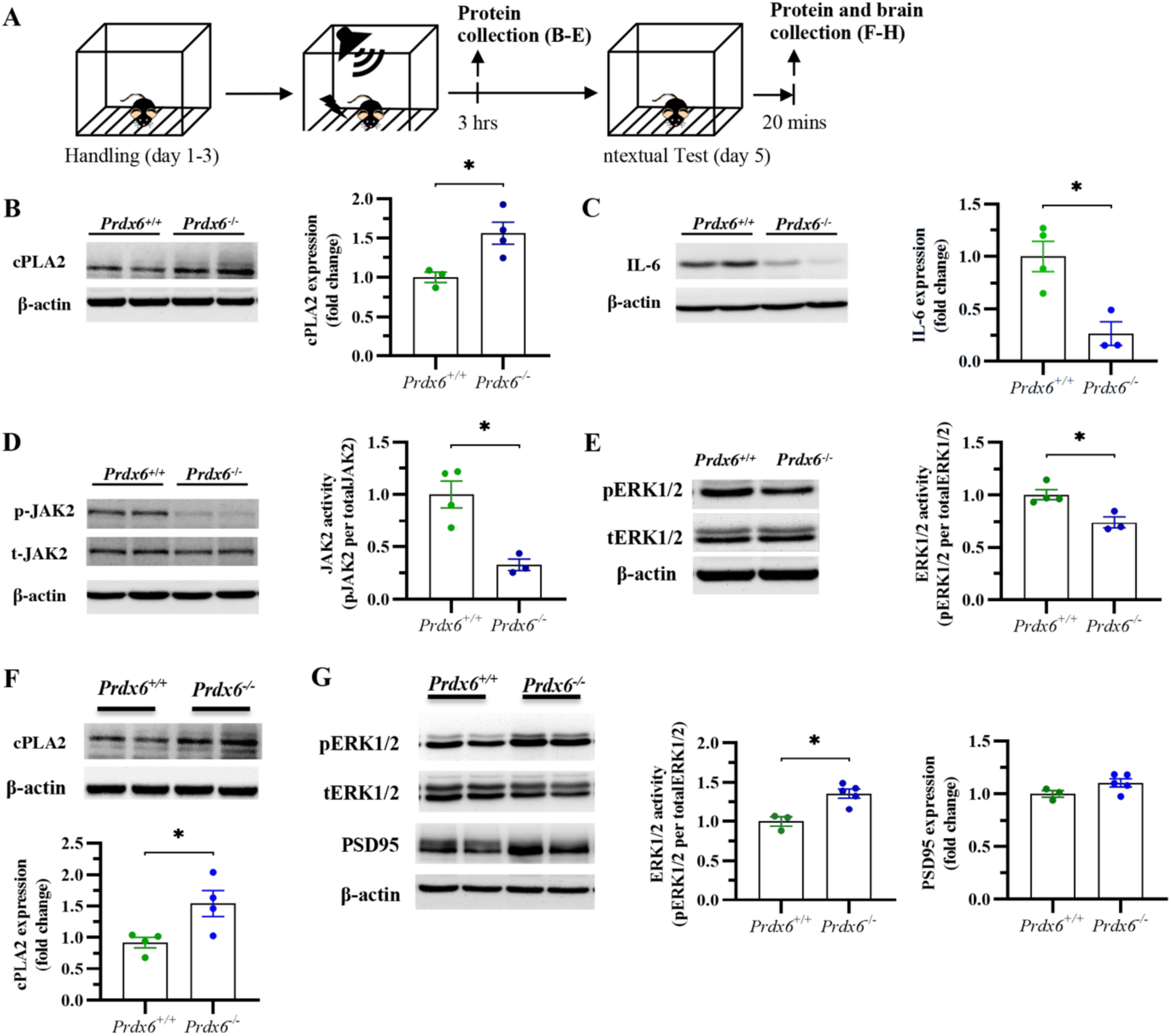
The expressional changes of memory-associated molecules during consolidation and retrieval of the contextual memory. **(A)** Hippocampal protein samples were collected 3 hours after training for consolidation process and 20 minutes after the contextual test for retrieval process. **(B-E; left)** Immunoblots of the expression of cPLA, IL-6, pJAK2, total JAK2, pERK1/2, total ERK1/2 and β-actin during memory consolidation. **(B-E; right)** Quantification data for the expression of cPLA2 (n = 3-4 per group, Student’s *t* test), IL-6, phosphorylated JAK2 and phosphorylated ERK1/2 in the hippocampus. **(F; upper panels)** Immunoblots of cPLA2 and β-actin expression in the hippocampus during memory retrieval. **(F; lower panels)** Quantification data of the expression of cPLA2 in the hippocampus of mice. **(G; left)** Immunoblots of pEKR1/2, total ERK1/2, PSD95 and β-actin expression in the hippocampus during memory retrieval. **(G; right)** Quantification data of the expression of phosphorylated ERK1/2 and PSD95 in the hippocampus of mice. All data represent the mean ± the SEM. \**p* < 0.05. cPLA2, cytosolic phospholipase A2; IL-6, interleukin 6; pJAK2, phosphorylated Janus kinase 2; tJAK2, total Janus kinase 2; PSD95, postsynaptic protein density 95.

Twenty minutes after the contextual test, the cPLA2 expression level was significantly increased in *Prdx6*^*−/−*^ mice (Fig. 6F) (*t*_6_ =−2.761, *p* = 0.033), and significantly higher ERK1/2 phosphorylation was recorded compared with those of their wild-type littermates (Fig. 6G) (*t*_5_ = −5.336, *p* = 0.003). However, no difference for postsynaptic density protein 95 (PSD95) was recorded (Fig. 6G) (*t*_*6*_ = −1.843, *p* = 0.115) in *Prdx6*^*−/−*^ mice.

### PRDX6 was localized in astrocytes and distributed in the CA1, CA3, and dentate gyrus of the hippocampus

We next investigated the localization and distribution of PRDX6 in the hippocampus after retrieval of contextual fear memory. Consistent with previous studies that showed high expression of PRDX6 in astrocytes (29), our results also revealed that PRDX6 was colocalized with the astrocytic marker, GFAP, in the CA1, CA3 and dentate gyrus of the hippocampi (Fig. 7A).

**Figure 7.**
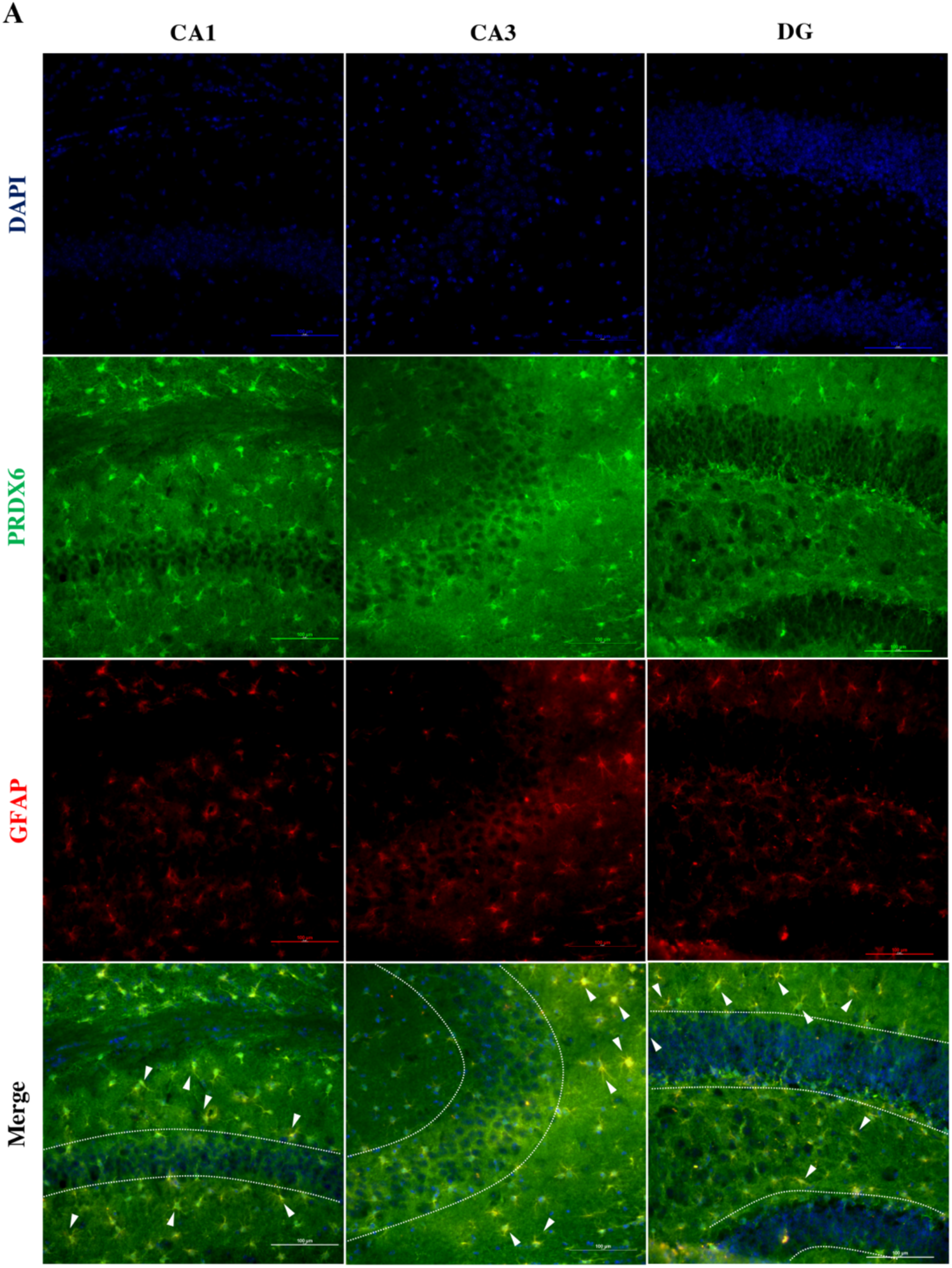
The expression of PRDX6 in hippocampal astrocytes after contextual testing. **(A)** Immunofluorescent images of brain sections from *Prdx6*^+*/*+^ mice after contextual testing. The merged images of double staining for PRDX6 (green) and GFAP (red) showing the colocalization (yellow, white arrows) of PRDX6 with GFAP in the CA1, CA3 and DG subregions of the hippocampus. PRDX6, peroxiredoxin 6; GFAP, glial fibrillary protein; CA1, cornu ammonis 1; CA3, cornu ammonis 3; DG, dentate gyrus.

## Discussion

The present study is the first report regarding the role of *PRDX6* in the regulation of fear response. Since the intraventricular injection of LV-hPRDX6-V5 was able to reverse the enhanced fear response of the *Prdx6*^−/−^ mice, we confirmed that the observed effect was attributable to PRDX6. Changes in several memory-associated molecules, such as cytosolic phospholipase A2 (cPLA2), interleukin 6 (IL-6), Janus kinase 2 (JAK2), and extracellular signal-regulated kinases 1 and 2 (ERK1/2), were also detected during memory formation and after memory retrieval.

Abnormal fear responses corresponding to symptoms of PTSD have been intensively investigated by using the fear conditioning paradigm (30, 31). The behavioral phenotypes of faster learning and enhanced fear memory retention after trace fear conditioning in *Prdx6*^*−/−*^ mice found in this study are similar to those observed in *Atf3* (32)and *Ccr5* (33) knockout mice. Using a computer-based search (Alggen Promo software, version 8.3, available on the web page: http://www.alggen.lsi.upc.es/cgi/bin/promo), we found that the promoter region of the *PRDX6* gene contains binding sites for activating transcription factor 3 (ATF3), suggesting the role of ATF3 in *Prdx6* transcription. Notably, knocking out of *Prdx6* resulted in a significant decrease in the C-C chemokine ligand 5 (CCL5) level, a ligand for CCR5 (34). These data indicate that ATF3, PRDX6 and CCL5/CCR5 may participate in the same or related pathways in the context of regulating fear memory performance.

Since we did not observe the change in reactive oxygen species (ROS) in *Prdx6*^*−/−*^ mice, we assume that the abolished peroxidase activity of PRDX6 may not play a dominant role in memory retrieval, at least in our study. On the other hand, deletion of aiPLA2 activity in *Prdx6*^*−/−*^ mice upregulated cPLA2 and increased ERK1/2 phosphorylation during memory retrieval stage may account for the *Prdx6* role in the regulation of fear response. Phospholipases A2 (PLA2s) are enzymes that cleave phospholipid to release arachidonic acid (AA) and lysophosphotidic acid (35). Both cPLA2 and iPLA2 can hydrolyze the sn-2 position of synaptic membrane phospholipids to generate AA and Docosahexaenoic acid (DHA) (36). AA and DHA are abundant in the brain cells and important for brain structure and functions (36, 37). The cPLA2 and phosphorylated ERK1/2 are also known to be important for memory retrieval (38-40). In the *Prdx6*^*−/−*^ mice, upregulation of cPLA2 may compensate loss of iPLA2 activity to maintain efficiency of synaptic membrane function, in turn, lead to the phenotype with enhanced fear memory.

In the hippocampal CA1, NMDA receptor-dependent long-term potentiation (LTP) is a form of synaptic plasticity and is a widely acceptable cellular mechanism of memory formation (41, 42). Enhanced memory performance is usually linked with increased LTP, however, here we found although the *Prdx6*^−/−^ mice exhibited enhanced fear memory, its late-phase LTP was reduced. Such exceptions were also reported by several previous studies (43, 44), and underlying mechanisms remained to be settled. Phosphorylation of ERK1/2 (pERK1/2) mitogen-activated protein kinase (MAPK) cascade is known to be required for the induction of early and late-LTP (45, 46). The decreased pERK1/2 detected in the *Prdx6*^−/−^ mice may be responsible for reduction of late-LTP.

The fact that PRDX6 expression was reduced after restraint stress (RST), trace fear conditioning (TFC), and glucocorticoid (GC) treatment confirmed its role in traumatic stress response. Studies have demonstrated close links between PRDX6, ATF3, and GC (47), the stress-suppressed PRDX6 expression phenomenon may be due to the negative regulation of ATF3 by GC. We also demonstrate that PRDX6 is highly expressed in hippocampal astrocytes, which supports previous findings in mice and humans (29, 48, 49). Stress-induced neurohormonal responses is known to cause various effects on neuronal activity through the activation of astrocytes, in turn, resulting in behavioral adaptation to cope stressful situations (50, 51). Further investigation by specifically inhibiting PRDX6 expression in astrocytes may help to elucidate the astrocyte-neuron interaction underlying traumatic stress response.

Taken together, our findings demonstrate that the ablation of *Prdx6* enhances freezing response to conditioned fear via the modulation of cPLA2, ERK1/2 and IL-6/JAK2 signaling in the hippocampus. Our results may help to understand the PTSD pathology and develop PRDX6 as a new therapeutic target.

## Materials and Methods

### Animals

All experiments on animal were approved by the Institutional Animal Care and Use Committee of Tzu Chi University, Taiwan (approval #104099), and comply with the Taiwan Ministry of Science and Technology guidelines for the ethical treatment of animals. Twelve-to 14-week-old wild-type (C57BL/6J) and *Prdx6*^*−/−*^ mice were originally generated by Wang X. (51) and provided by Dr. Shun-Ping Huang at Tzu Chi University, Taiwan. All mice were maintained in the Laboratory Animal Center of Tzu Chi University and were housed with *ad libitum* access to food and water under a constant 12-hour light/dark cycle. Genotyping (Fig. S1A), qRT-PCR (Fig. S1B) and western blot analysis (Fig. S1C) were conducted to confirm the absence of *Prdx6* in knock-out mice before every behavioral test. Moreover, we also recorded the morphology and body weight of the *Prdx6*^*−/−*^ mice. We found that both morphology (Fig. S1D) and body weight (Fig. S1E) [*t*_19_ = −1.426, *p* = 0.170] of the *Prdx6*^*−/−*^ mice appeared to be normal.

### Behavioral experiments

#### Trace fear conditioning

Trace fear conditioning was modified from the protocol used in our previous study (30). The conditioned chamber (17 cm (W) x 17 cm (L) x 25 cm (H)) illuminated with a white 30-lux light under the top-view camera was used in this study. After three days of habituation, mice were trained with three pairs of tone (CS) and electric foot shock (US) with an inter-trial interval of 1 min for a total of 9 min. One pair of CS-US consisted of 20 s tone (6000 Hz, 85 dB) and 1 s electric foot shock (2 mA) with 10 s training interval. To test their contextual fear memory retention, the mice were re-exposed to a conditioned chamber for 6 min without giving any tone and foot shock after 24 hours of training session. The freezing behavior, defined as no movement except breathing was analyzed using tracking software (EthoVision XT 15, Noldus Information Technology). The freezing time was converted to freezing percentage using the following formula:

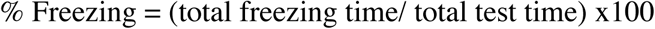

### Restraint stress (RST)

To mimic PTSD model, mice in RST group were received 30 min of immobilization in 50 ml plastic conical tubes with breathing holes covered by non-transparent paper box to mimic the dark phase (17). Immediately after 30 min of RST, stress mice were sacrificed immediately for protein collection.

#### Open field test

An open chamber (50 cm (W) x 50 cm (L) x 50 cm (H)) was used to test locomotor function of the mice under light on condition. The movements of the animals were recorded by the camera hung on top and their locomotor activity along 10 min of the test. The movements of the animals were recorded by the camera hung on top and their locomotor activity (distance travelled and moving speed) and time spent were measured and analyzed by the tracking software (EthoVision XT 15, Noldus Information Technology).

#### Elevated-Plus Maze Test

The elevated-plus maze is used to assess the anxiety-related behavior in rodents. The apparatus consists a “plus”-shaped maze at 60 cm height above ground with two oppositely positioned closed arms and two oppositely positioned open arms and a centre region. The mice were placed in the centre region facing one of the closed arms and allowed to explore the maze freely for 10 minutes. A video camera and tracking system (EthoVision XT 15, Noldus Information Technology) were used to record and analyze their anxiety-like behavior, respectively.

### Lentiviral vector preparation for PRDX6

Total RNA was isolated from ARPE cells and converted to cDNA using oligo (dT) 18 primers. The cDNA was then amplified using a specific forward primer (5’-ATG CCC GGA GGG TTG CTT C-3’ containing an XbaI site) and reverse primer (5’-GC GAA TTC TTA AGG CTG GGG TGT ATA ACG-3’containing an EcoRI site) (52). Full-length human Prdx6 cDNA was purified by a PrestoTM Mini Plasmid Kit (catalog #, Geneaid Biotech Ltd., Taiwan). To add the V5 tag to the C-terminus of PRDX6, the purified PCR products and pcDNA4/V5-His A vector (catalog #, Introgen) were cut with XbaI and EcoRI. pLAS3wPpuro vectors containing EGFP and PRDX6-V5 were designed for the production of lentiviral vectors. HEK293T cell lines were used to produce lentiviruses containing the EGFP and PRDX6-V5 genes. Harvested lentivirus was concentrated using PEG-it (™) virus precipitation solution (System Biosciences, CA) and processed for titration.

### Stereotaxic surgery and intralateral ventricular injections of lentivirus containing human PRDX6-V5

The procedures for stereotaxic injection were performed according to our previous study (53) with slight modification. Briefly, mice were anesthetized by intraperitoneal (IP) injection of ketamine/xylazine mixture (0.45 ml/ 25 g of body weight) and then fixed on the stereotaxic frame (Stoelting, US). The lentivirus containing either EGFP or human PRDX6-V5 was dissolved in sterile 1X phosphate-buffered saline (PBS) to obtain the final titer of 7 ×10^5^ in 2 µl volume. The lentiviral vectors were then unilaterally injected into the right lateral third ventricle with the following brain coordinates: anterior posterior (AP), −0.5 mm; medial lateral (ML), 1 mm (from the bregma): and DV, 2.33 mm (from the skull surface). A 10-ul Hamilton syringe with a 26 G needle was placed on the microinfusion pump (KD Scientific Inc. MA, USA) and connected via polyethylene-28 mm I.D. tubing to the internal cannula. The lentiviruses were injected with a flow rate of 0.5 µl/min over a period of 4 min. The cannulas were placed for another 5 min to allow diffusion before removing them. Following surgery, mice were given pain killers (meloxicam, Achefree, Taiwan) and allowed to recover for 2.5 weeks before the behavioral tests.

### Cell culture

A Human Retinal Pigment Epithelial Cell Line (ARPE-19) were cultured in Dulbecco’s Modified Eagle Medium: nutrient mixture F-12 (DMEM/F12) containing 10% fetal bovine serum, 100U/ml penicillin, and 100ug/ml streptomycin. The cells were incubated at 37°C in a humidified 5% CO_2_ atmosphere. The cells were grown until 80% confluence and used at passage 20-23.

### Cell viability assay

Cells were plated in 96-well plates for 24 hours. Then cells were treated with 0.01% DMSO or varying doses (1, 10 and 100 nM) of glucocorticoid (GC) for 1 hour. Cell viability was measured by Thiazolyl Blue Tetrazolium Blue (MTT). Briefly, 10 µl of MTT solution (5mg/ml) was added to each well and incubated for 3 hours at 37°C. After removing the supernatant, 100 µl DMSO was added in each well. The intensity is measured calorimetrically at 570 nm with a microplate reader (Thermo Scientific Multiskan Spectrum, USA).

### Extracellular LTP recording

The 12 to 14-weeks old *Prdx6*^+*/*+^and *Prdx6*^*−/−*^ mice were anesthetized with isoflurane and sacrificed by guillotine. Whole brains were immediately removed and washed in ice-cold artificial cerebrospinal fluid (ACSF). Immediately, the hippocampi were extracted and placed in the groove of a 3% agarose block for sectioning. The posterior 1/5 of the hippocampus was trimmed giving a flat edge that will then help to stabilize the hippocampi on the metal chamber of a vibrating microtome (Microslicer DTK-1000, Dosaka EM Co. Ltd., Kyoto, Japan). The hippocampi were horizontally sectioned in oxygenated ACSF at 450 μm thickness. The hippocampal slices were transferred and remained on the filter paper containing ACSF in the incubated boxes at room temperature (27 °C) for 90–120 min. The hippocampal slices were transferred to a recording chamber for extracellular LTP recording. During recording, slices were perfused with ACSF containing 0.1 mM picrotoxin at a speed of 2–3 mL/min. The glass pipettes were pulled on a micropipette puller (PC-10 Needle Puller, Narishige, Japan) filled with 3 M NaCl. This recording electrode was placed at the stratum radiatum of the CA1 for recording the excitatory postsynaptic field potentials (fEPSPs). The bipolar stainless-steel microelectrodes (Frederick Haer Company, Bowdoinham, ME, USA) were used as a stimulus electrode. The stimulation intensity was adjusted between 0–80 μA for each slice, so that the fEPSPs were elicited to approximately 50% of the maximal response. Baseline fEPSPs were evoked every 15 s for 20 min followed by high-frequency stimulation (HFS), which includes 3 trials of 100 pulses at 100 Hz for 60 s. Then, fEPSPs were stimulated every 15 s for an additional 60 min. Recordings were amplified using an Axon Multiclamp 700B amplifier. All signals were filtered at 1 kHz and digitized at 10 kHz by CED Micro Power 1401 mKII interface (Cambridge Electronic Design, Cambridge, UK) using Signal software. The downward slope of fEPSPs was measured.

### Immunofluorescence staining

For immunocytochemistry, cells were plated on cover slides and grown in 24-well plates. After 1 hour of GC treatment, cells were washed with 1X PBS three times and then fixed with 4% paraformaldehyde for 30 minutes at room temperature. For immunohistochemistry, mice were anesthetized and transcardially perfused using 0.9% saline and 4% paraformaldehyde. Brains were exercised immediately and post fixed with 4% PFA for another 2 days. After that, the brains were washed with 1x PBS three times and then stored in 30% sucrose at 4°C. After the dehydration period, the brains were embedded in optimal cutting temperature compound (Sakura Finetek USA, Inc., USA) and stored at −80°C until sectioning. Cryopreserved brains were sectioned at 20 µm using cryostat. The staining protocol for both cells and tissues are the same in which samples were washed with a washing buffer (1x PBS containing 0.3% Triton X-100) and further blocked with 1 mg/mL bovine serum albumin containing 0.3% Triton X-100 for 1hr. Subsequently, samples were double-stained with monoclonal goat anti-GFAP (1:200, Abcam, UK) and polyclonal rabbit anti-PRDX6 (1:200, Bethyl laboratories, Inc, USA). Then the samples were washed with washing buffer and incubated in secondary antibody: Alexa Fluor 546 anti-goat and Alexa Fluor 488 anti-rabbit IgG (1:200, ThermoFisher Scientific, USA) for 1hr, followed by washes with PBS, and counterstained with DAPI (1:10,000) for 5min. The images were obtained by fluorescent microscope (Nikon model# ECLIOSE Ni-E, Japan)

### Detection of oxidative stress levels in the hippocampus

To measure ROS levels in the hippocampus, mice were sacrificed and the brains were isolated 20 min after contextual test. Briefly, the fixed brains were sectioned by cryostat with 20 µm thickness. Hippocampal sections were then immersed in 1 μmol/L dihydroethidium (DHE) in PBS solution at room temperature for 5 minutes. The stained sections were washed with 1X PBS three times, and cover-slipped. DHE is oxidized by superoxide anion to form ethidium binding to DNA in the nucleus and emits red fluorescence. The imaging procedure was done under fluorescent microscope (Nikon model #ECLIOSE Ni-E, Japan) with excitation /emission wavelength of 380/420 nm.

### Western blotting

The mice were sacrificed immediately after the completion of restraint stress. Under trace fear conditioning, hippocampal proteins were extracted at 3 hours after training and 20 minutes after the contextual test. After decapitation, the hippocampi were isolated and homogenized in ice cold RIPA lysis buffer 1X (Millipore, USA) containing protease and phosphatase inhibitor. The protein samples were kept on ice for 30 min before centrifuge at 13,000 rpm for 15 min at 4°C. The supernatants were collected for further experiments. For non-reducing SDS-PAGE, protein (30 or 45 ug) samples were boiled at 95°C in 1X sample buffer without reducing agent for 10 minutes and samples were cooled for 5 mins. Similar to non-reducing condition, except adding reducing agent into protein samples was included to study total PRDX6 and other proteins of interest under reducing condition. The samples were loaded and run on 8% or 10% SDS-PAGE at 80 mV in stacking gel and 120 mV in resolving gel. The separated proteins were then transferred to a PVDF membrane (0.2 and 0.4 um pore size). The blots were incubated with anti-cPLA2 (1:1000; Santa Cruz, USA), anti-IL6 (1:2000; Proteintech, USA), anti-pJAK2 (1:500; Cell Signaling, Danvers, MA), anti-JAK2 (1:1000; Cell Signaling, Danvers, MA), anti-pERK1/2 (1:1000; Cell Signaling, Danvers, MA), anti-total ERK1/2 (1:1000; Cell Signaling, Danvers, MA), anti-PSD95 (1:2000; ThermoFisher, USA), anti-PRDX6 (1:2000; Abcam, UK) or anti-β-actin antibody (1:10000; Sigma-aldrich) in TBST containing 0.1% BSA (ThermoFisher, USA) overnight at 4°C room on a shaker. On the next day, the blots were incubated with horseradish peroxidase-conjugated secondary antibody goat anti-mouse IgG (Cell signaling, Danvers, MA) for cPLA2, PSD95, PRDX6, and β-actin and goat anti-rabbit (Santa Cruz Biotechnology, Santa Cruz, CA, USA) for IL-6, pJAK2, JAK2, pERK1/2 and ERK1/2 with the dilution of 1: 10000 in blocking buffer for 1 hour at room temperature. After three washes for 5-min in TBST buffer, the membranes were developed using ECL (Western lightning ® Plus ECL, PerkinElmer Inc, MA, USA) and detected under UVP Biospectrum 810 imaging system. The band intensities were quantified using ImageJ 1.52a (National Institutes of Health, USA).

### Data analyses

Statistical analysis was performed using SPSS (version 25, IBM Corporation), and the graphs were made using GraphPad Prism version 8. All data are presented as mean ± SEM, with statistically significant at *p* < 0.05. Student’s *t* tests were conducted when compared to two independent groups. For multiple comparisons, one-way ANOVA followed by Bonferroni post hoc analysis was used. For related, not independent groups were analyzed by two-way repeated measures ANOVA. Sample sizes are indicated in figure legends.

## Acknowledgments

This work was supported by the Ministry of Science and Technology (MOST), Taiwan (MOST-107-2410-H320-DOI-MY3), and Tzu Chi University/Tzu Chi Foundation (TCMF-SP-108-04).

## Competing interests

The authors report no biomedical financial interests or potential conflicts of interest.

---

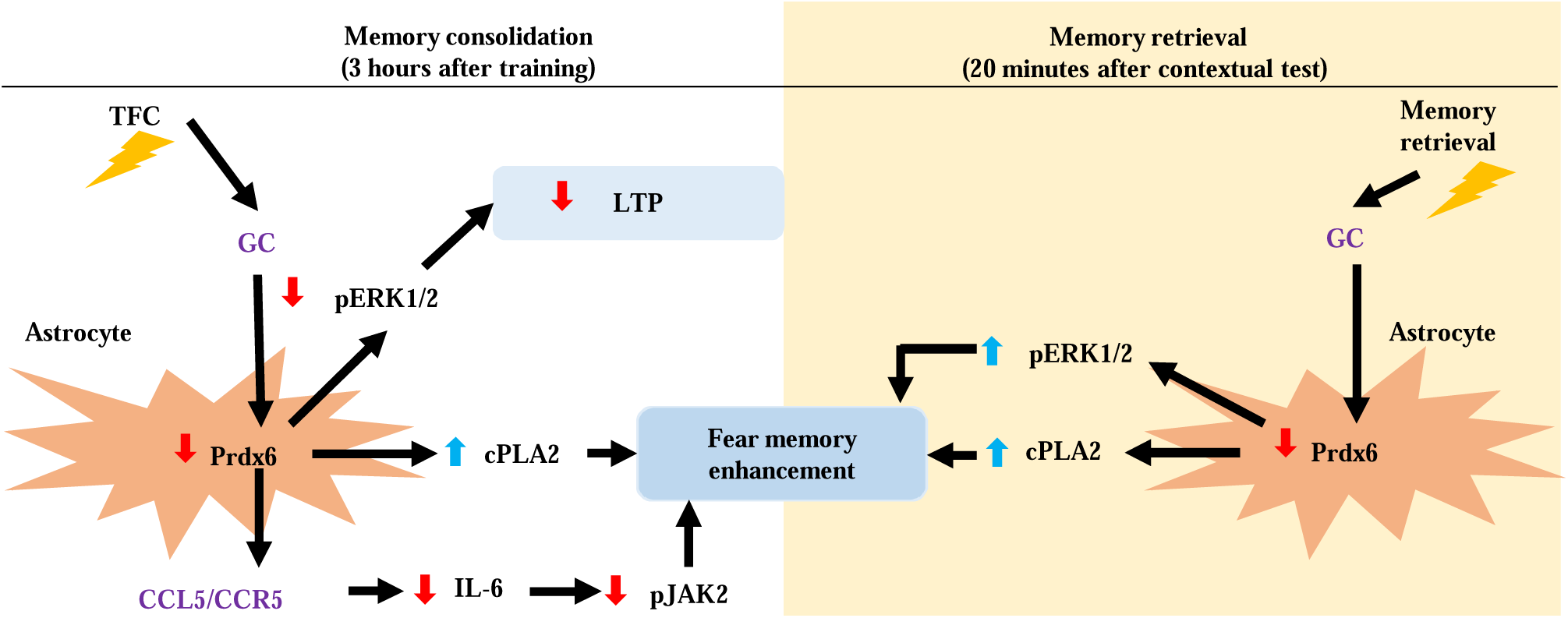

## Notes

### Competing Interest Statement

The authors have declared no competing interest.

